# Activity of FoxP2-positive neurons is associated with tadpole begging behavior

**DOI:** 10.1101/2023.05.26.542531

**Authors:** Sarah C. Ludington, Jordan E. McKinney, Julie M. Butler, Billie C. Goolsby, Ashlyn A. Callan, Maiah Gaines-Richardson, Lauren A. O’Connell

## Abstract

Motor function is a critical aspect of social behavior in a wide range of taxa. The transcription factor FoxP2 is well studied in the context of vocal communication in humans, mice, and songbirds, but its role in regulating social behavior in other vertebrate taxa is unclear. We examined the distribution and activity of FoxP2-positive neurons in tadpoles of the mimic poison frog (*Ranitomeya imitator*). In this species, tadpoles are reared in isolated plant nurseries and are aggressive to other tadpoles. Mothers provide unfertilized egg meals to tadpoles that perform a begging display by vigorously vibrating back and forth. We found that FoxP2 is widely distributed in the tadpole brain and parallels the brain distribution in mammals, birds, and fishes. We then tested the hypothesis that FoxP2-positive neurons would have differential activity levels in begging or aggression contexts compared to non-social controls. We found that FoxP2-positive neurons showed increased activation in the striatum and cerebellum during begging and in the nucleus accumbens during aggression. Overall, these findings lay a foundation for testing the hypothesis that FoxP2 has a generalizable role in social behavior beyond vocal communication across terrestrial vertebrates.

## Introduction

In species where parents provision their young, offspring signaling can be important for obtaining food. Begging behavior generally involves coordination of motor circuits, such as vocalization in chicks [1–3], vibrational displays in amphibian tadpoles [4,5], and chemical and motor signals in insect larvae [6,7]. There is a rich theoretical literature on the evolution of offspring signaling and parental investment [8,9], which has been experimentally investigated mostly in birds [10–12]. While the behavioral and physiological ecology of begging has received much attention, the neural basis of this critical behavior is relatively unknown, with the exception that bird begging behavior uses the same vocal-motor pathways later used in adult song [13]. Investigating the neural circuits and gene networks that regulate offspring signaling would establish a mechanistic view on how begging behavior evolves from ancestral neural features. Additionally, this perspective would complement the existing theoretical models of how and when begging signals evolve.

Forkhead Box P2 protein (FoxP2) is associated with motor processes related to behavior in many species. This protein is a transcription factor that regulates gene networks important in many neuronal functions, including genes involved in synaptic plasticity, neurotransmission, and axonal guidance [14,15]. Interest in FOXP2 surged when a mutation was linked to speech and language impairments in humans [16] (human FOXP2 and non-human FoxP2 homologs are upper and lowercase, respectively). The mutation of a critical residue in the DNA-binding domain of the human FOXP2 (the R553H mutation) causes difficulty with fine rapid movements of the mouth and face that impair speech [17]. FOXP2 truncations and intragenic deletions also manifest in language and speech impairments [18–20]. Individuals carrying FOXP2 disruptions are at risk for other phenotypes such as difficulties feeding in infancy and low performance in receptive and expressive language assessments [21]. A conserved role for FoxP2 has been extended to other species, where FoxP2 manipulations influence vocalizations emitted by birds [22–24] and mice [25–27]. Functional studies in humans, mice, and birds point to the role of FoxP2 in the development and function of corticostriatal and cortico-cerebellar circuits important for motor control [16,28,29]. Despite the research emphasis on vocal communication, FoxP2 manipulations in mice lead to altered social interactions, like parental care and aggression [30,31], as well as skilled motor tasks [32], suggesting a broad role for FoxP2 in coordinating motor aspects of behavior.

Frogs use both vocal and non-vocal signaling for social interactions [33,34], but the role of FoxP2 in frog communication has not yet been investigated, to our knowledge. Like in mammals [35], FoxP2 is expressed in early brain development of *Xenopus* [36]. Given expression of FoxP2 in the larval brain and that FoxP2 is important in regulating motor aspects of social behavior in mammals and birds, we reasoned that FoxP2 may play a role in amphibian social behaviors as well. Specifically, since FoxP2 has an important role in vocal-motor pathways of bird song [22–24], which are more active in begging birds [13], we reasoned that FoxP2 may be involved in tadpole begging behavior. Additionally, as FoxP2 also leads to altered aggression in mice [30], we reasoned that FoxP2 may also be associated with tadpole aggression. In the present study, we tested the hypothesis that FoxP2 is associated with begging signals of tadpoles towards adult conspecifics or aggression towards conspecifics. We tested this hypothesis in the mimic poison frog (*Ranitomeya imitator*), where tadpoles beg parents for unfertilized egg meals by vigorously vibrating back and forth with their heads and nipping at visiting females with their mouths [37]. Tadpoles are reared in isolated nurseries where they are aggressive to intruder tadpoles [4,38]. We first mapped the neural distribution of FoxP2 in the *R. imitator* tadpole brain and then compared the activity of FoxP2-positive neurons across begging, aggressive, and control animals. Given the extensive literature of striatal FoxP2 in vocal communication in birds and mice, we predicted that FoxP2-positive neurons in the striatum would have higher activity in begging tadpoles.

## Methods

### Animals

*Ranitomeya imitator* tadpoles were bred from our laboratory colony [39]. Adult *R. imitator* females from breeding pairs were used as stimulus animals in the begging context. A conspecific tadpole was used as a stimulus in the aggression context. All procedures were approved by the Stanford University Animal Care and Use Committee (Protocol #33097).

### Behavior

We randomly assigned tadpoles (Gosner stage 30-34, no forelimb development and minimal hindlimb development) into one of three experimental groups: a reproductive adult female (begging, N=14); a smaller sized conspecific tadpole (aggression, N=15), or exposed to a novel object (a metal bolt, N=15). Conspecific stimuli were different between trials. All behavior trials were conducted between 09:00 and 12:00 hours. Tadpoles were placed into individual square arenas (5 × 5 × 5 cm) filled with 50 mL of conditioned water (Josh’s Frogs R/O Rx, Owosso, MI). Tadpoles were recorded from above using GoPro cameras (GoPro HERO7 Black, 1080p, 240 fps). Each tadpole acclimated for 10 min in the arena. Then, the stimulus was introduced to the arena and behavior was recorded for 30 min. Stimuli were then removed from the arena and tadpoles were placed in the dark for 15 minutes to minimize post stimulus neural activity. This additional time was included as pilot experiments with pS6-immunoreactivity suggests this marker peaks 45 min post stimulus. Tadpoles were then anesthetized with topical 20% benzocaine and euthanized by decapitation.

Videos were scored using BORIS software [40] by an observer uninformed of tadpole identity (Figure S1). Begging was quantified by the number and duration of each begging bout, where the tadpole orients to, intensely vibrates near, and occasionally nips at a conspecific. Aggression and cannibalism is observed in tadpoles of this species, where tadpoles will attack and consume conspecifics. In this study, aggression was quantified by the number and duration of attacks towards the other tadpole. Control tadpoles did not display either of these behaviors.

### Immunohistochemistry

Whole tadpole heads were fixed with 4% paraformaldehyde (PFA) in 1X phosphate buffered saline (PBS) at 4°C overnight, rinsed in 1X PBS, and transferred to a 30% sucrose solution for cryoprotection at 4°C overnight. Samples were then embedded in mounting media (Tissue-Tek® O.C.T. Compound, Electron Microscopy Sciences, Hatfield, PA, USA) and stored at -80°C until cryosectioning at 15 μm into three series. Sections were thaw-mounted onto SuperFrost Plus microscope slides (VWR International, Randor, PA, USA) and then stored at -80°C until immunohistochemistry.

We used double-label fluorescence immunohistochemistry to detect FoxP2 and phosphorylated ribosomes (pS6, phospho-S6) as a proxy of neural activity [41], as previously described [42]. Slides were incubated overnight in a mix of both primary antibodies [rabbit anti-pS6 (Invitrogen, cat #44-923G) at 1:500 and goat anti-FoxP2 (Abcam, cat #AB1307) at 1:500 in 2% normal donkey, 0.3% TritonX-100, 1X PBS]. Following several washes, slides were incubated in a mix of fluorescent secondary antibodies (1:200 Alexa 488 donkey anti-goat and 1:200 Alexa 568 donkey anti-rabbit in 2% normal donkey serum, 0.3% TritonX-100, 1X PBS) for two hours. Slides were then rinsed in water and cover slipped using Vectashield Hardset Mounting Medium with DAPI (Vector Laboratories, Burlingame, CA, USA) and stored at 4°C. FoxP2 was restricted to cell nuclei and additional antibody characterization can be found in Supplementary Materials (Figure S2-S5).

### Fluorescence microscopy and cell counting

Brain sections were imaged on a Leica compound fluorescent microscope with a QImaging Retiga 2000R camera as previously described [42]. Brain regions containing FoxP2 were identified using DAPI-stained nuclei while referencing a poison frog brain atlas [42]. FIJI software [43] was used to measure the area of the nucleus accumbens, striatum, and cerebellum within a single hemisphere. The number of FoxP2-positive cells, pS6-positive cells, and colocalized cells were quantified within each area using the “Cell Counter” function. Due to tissue quality, one to four sections were counted per individual per brain region.

### Data analysis

All statistics and figures were generated in R Studio (version 1.1.442) running R (version 3.5.2). We used the glmmTMB R package [44] to analyze cell count data with generalized linear mixed models. For FoxP2-positive and pS6-FoxP2 colocalized cells, we ran separate models using a negative binomial distribution; model fit was confirmed using DHARMa [45]. For both models, we tested the main effects of the experimental group (begging, aggression, control), brain region, and their interaction. Tadpole identity was included as a random variable to account for repeated sampling of brain regions within individuals. The log of the brain region area was included as an offset. For colocalization data, the number of colocalized (pS6 + FoxP2) cells was the dependent variable and the number of FoxP2 cells was included as a weight in the model. We then used the Anova.glmmTMB function for reported statistical values. When there was a significant interaction between group and brain region, we ran a post-hoc test with the emmeans R package (version 1.5.3) and used false discovery rate correction for multiple hypothesis testing. Correlations between behavior and cell counts were tested using the cor.test function in the R base package with the Spearman method.

## Results

### Neural distribution of FoxP2

We observed a broad distribution of FoxP2-positive cells throughout the tadpole brain (Figure 1, Figure S6). The highest densities of FoxP2-positive cells were found in the subpallial forebrain, optic tectum, thalamus, and cerebellum. Notably, there were many FoxP2 cells in regions linked to sensory processing, such as the olfactory bulb (chemosensory), torus semicircularis (acoustic processing), and optic tectum (vision).

**Figure 1.**
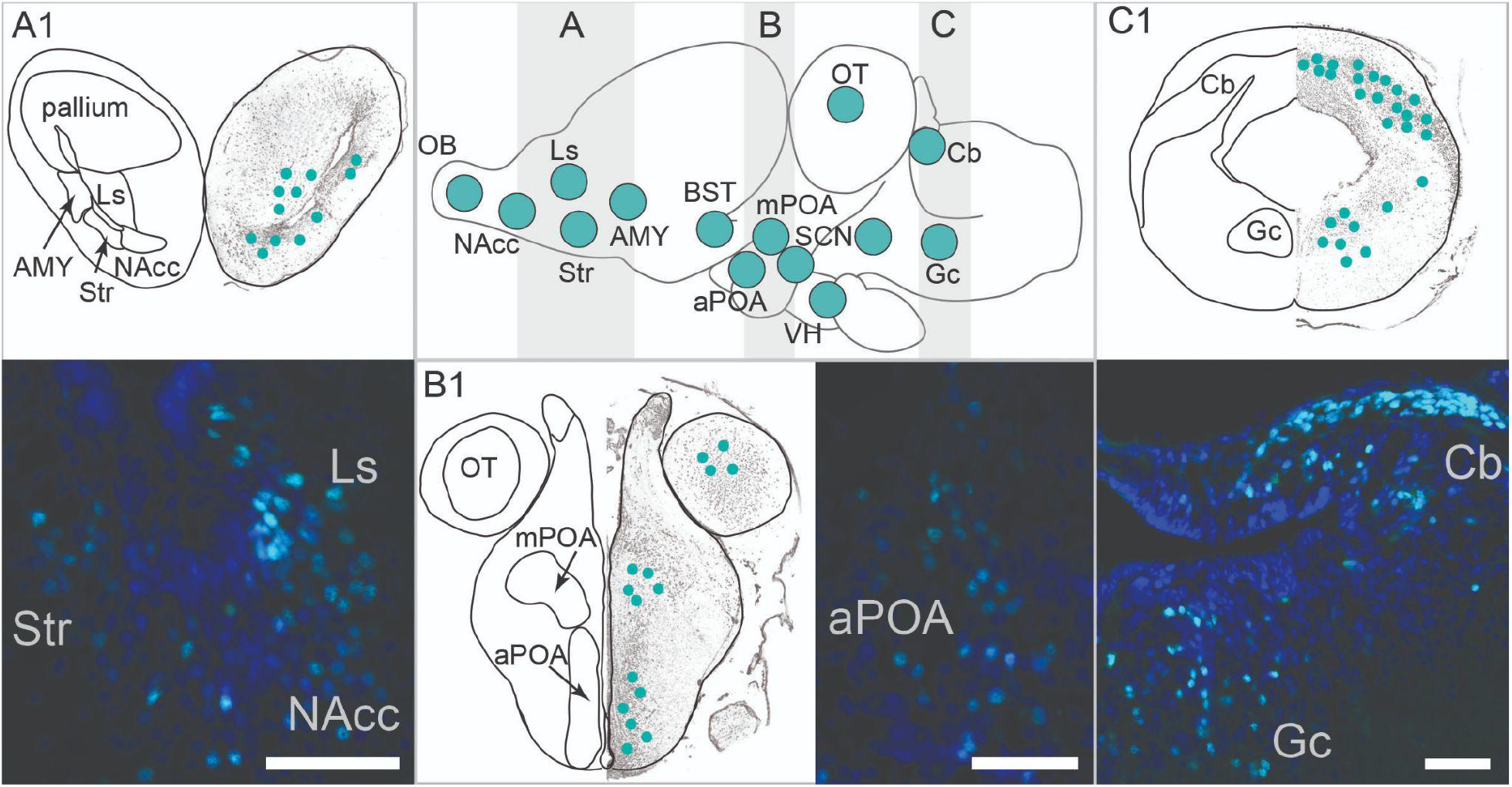
Neural distribution of FoxP2 in amphibians is similar to other vertebrates. FoxP2 is widely distributed throughout the amphibian brain, including the subpallial forebrain (A), midbrain (B), and a few hindbrain regions (C). The center sagittal brain schematic (rostral is to the left) shows brain regions (green) with FoxP2 positive cells. Gray boxes represent areas of interest for more detailed neuroanatomy and micrographs (A1-C1), where green dots represent qualitative presence of FoxP2. Micrographs show FoxP2-positive cells (cyan) and DAPI-stained nuclei (blue); scale bar is 20 μm. The complete neural distribution for FoxP2 can be found in Supplementary Materials. Abbreviations: AMY, amygdala; aPOA, anterior preoptic area; BST, bed nucleus of the stria terminalis; Cb, cerebellum; Gc, central gray; Ls, lateral septum; mPOA, magnocellular preoptic area; NAcc, nucleus accumbens; OB, olfactory bulb; OT, optic tectum; SCN, suprachiasmatic nucleus; Str, striatum; VH ventral hypothalamus.

### FoxP2-positive neuronal activity changes with different social stimuli

We investigated whether FoxP2-positive neuronal activity is associated with social behavior by quantifying the proportion of FoxP2-positive cells that colocalized with the pS6 marker of neural activity in tadpoles showing begging, aggression, or exposed to a novel object (asocial control) (Figure 2). We focused our quantification efforts on the basal ganglia (nucleus accumbens and striatum) and cerebellum given their robust expression of FoxP2 in mice and birds [46] and functional studies suggesting FoxP2-associated vocalization deficits are due to altered corticostriatal and corticocerebellar circuits [16,28,29]. The activity of FoxP2-positive cells depended on an interaction of behavioral group and brain region (group*region: F_4_ = 130.66, p < 0.001). Begging tadpoles had more active FoxP2-positive cells than aggressive and control tadpoles in the striatum (Str, aggression vs begging: z = -3.144, p = 0.005; begging vs control: z = 2.517, p = 0.018) and cerebellum (Cb, aggression vs begging: z = -2.490, p = 0.019; begging vs control: z = 3.626, p < 0.001). The number of active FoxP2-positive cells did not differ between aggressive and control animals in the striatum (p = 0.175) or cerebellum (p = 0.920). Aggressive tadpoles had more active FoxP2-positive cells than control tadpoles in the nucleus accumbens (NAcc, z = 2.989, p = 0.008), whereas activity of FoxP2-positive cells in this brain region did not differ between begging and aggression (p = 0.125) or control (p = 0.125) contexts. There was a significant difference in the number of FoxP2-positive cells within these brain regions across groups, where aggressive tadpoles had more FoxP2-positive cells in the striatum (Figure S7). There were no significant differences in the number of pS6 cells within these brain regions across groups and there were no significant correlations between activity of FoxP2-positive cells and measures of begging or aggressive behaviors (Figure S8).

**Figure 2.**
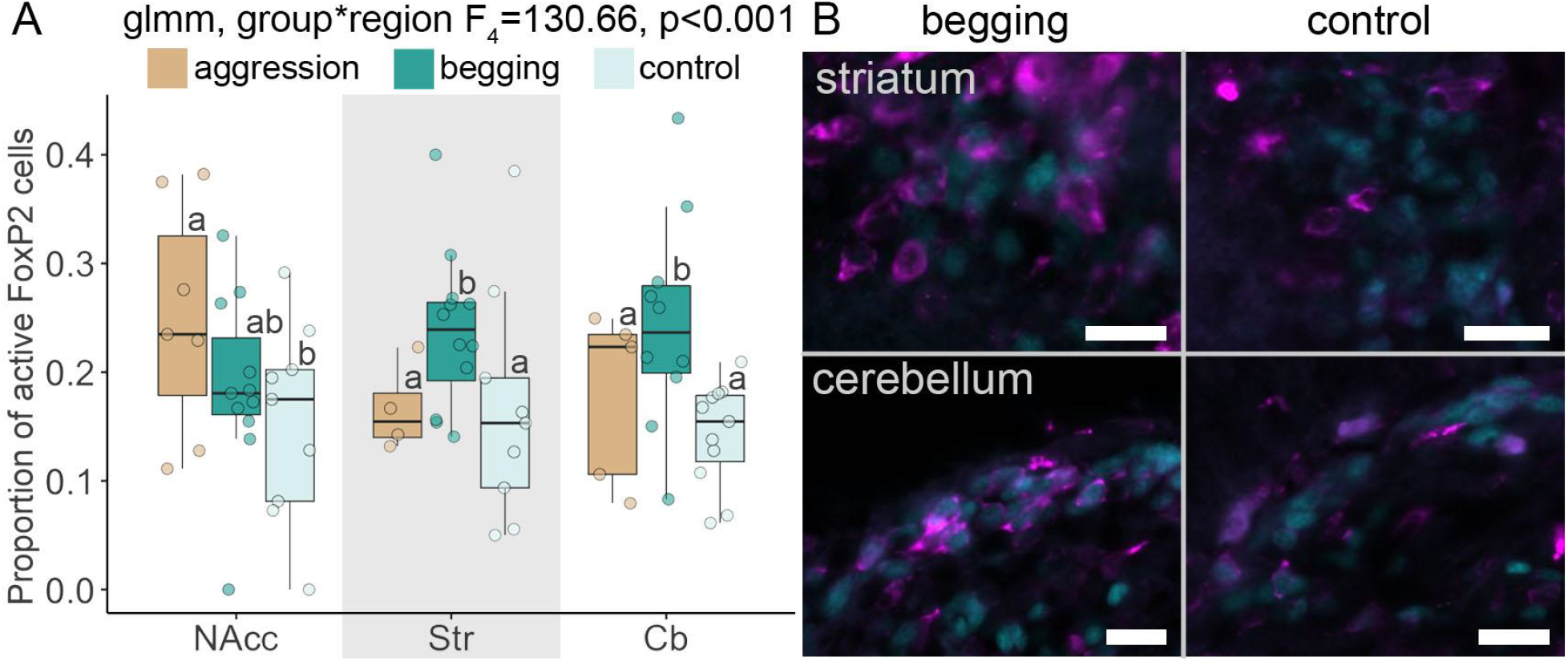
Activity of FoxP2 neurons changes with social behavior. **(A)** Proportion of active FoxP2-positive cells in aggressive (orange), begging (dark green) and control (light green) tadpoles are shown in boxplots with individual tadpoles displayed in dots. Within each brain region, groups not connected by the same letter are significantly different. **(B)** Representative micrographs of FoxP2 (green) and pS6 (pink) colocalization expression of begging (left) or control (right) tadpoles in the striatum and cerebellum. Scale bar is 10 microns. Abbreviations: Cb, cerebellum; NAcc, nucleus accumbens; Str, striatum.

## Discussion

Across species, there is variation in whether and how young animals display aggression or signaling to caregivers, but both behaviors require motor function coordinated by neural processes. Among other functions, the transcription factor FoxP2 plays a well established role in coordinating social behaviors like vocal communication in mammals and birds [17,20,24,28,47,48]. Our study expands the role of FoxP2 to social behavior in amphibians, laying a foundation for testing the generalizable function of FoxP2 in coordinating aspects of social behavior across terrestrial vertebrates in future studies.

### The brain distribution of FoxP2 is conserved across vertebrates

FoxP2 is widespread throughout the amphibian brain, with a distribution pattern consistent with those found in other vertebrates (mammals: [35,49,50]; birds: [51]; fish: [52–54]). Across these taxa, there is a conserved pattern of expression in brain areas involved in motor output, sensory processing, and sensorimotor integration. In *R. imitator* tadpoles, brain regions that regulate motor output and social behavior, including the basal ganglia and cerebellum, had many FoxP2-positive cells. FoxP2 is expressed in the basal ganglia and cerebellum in avian vocal and non-vocal learners, crocodiles, and rodents, suggesting conserved expression in motor-related areas regardless of the ability to learn acoustic communication [49,51]. Although there is conservation in the presence of FoxP2, its abundance is variable in songbirds depending on age and environment, suggesting FoxP2 expression may be linked to periods of vocal plasticity [51]. We also noted FoxP2-positive cells in many sensory processing regions like the olfactory bulb (chemosensory), optic tectum (visual processing), and torus semicircularis (acoustic processing). In bats, species differences in FoxP2 expression in the olfactory bulb are associated with different feeding habits (frugivorous versus insectivorous) [55], suggesting that FoxP2 may influence olfactory processing. Given *R. imitator* tadpoles rely on smell to distinguish between heterospecific stimuli [39], investigating FoxP2’s role in sensory integration broadly may be a valuable future research direction. However, the expression pattern of FoxP2 is variable across sex, age, and species [49,56], including species differences in expression of FoxP2 in neuronal and non-neuronal cells in the mammalian cortex [57]. This variability in expression makes the regulatory role of FoxP2 in behavior unclear, but a fruitful avenue of future research. Overall, the distribution of FoxP2 in the amphibian brain suggests a largely conserved pattern across terrestrial vertebrates.

### A general role for FoxP2 in social behavior

We found that activity of FoxP2-positive neurons was higher in the striatum and cerebellum of begging tadpoles and in the nucleus accumbens of aggressive tadpoles. Whether this pattern is directly relevant to these behaviors requires functional manipulations in a brain region-specific manner. Regardless, the context-dependent neuronal activation points to brain region specific roles for FoxP2-positive cells in social behavior. These findings lay a foundation for testing the hypothesis that FoxP2 has a generalizable role in social behavior beyond vocal communication.

The striatum is important for motor skills in many vertebrates [58] and has been linked to vocal communication in several taxa [59]. We found that FoxP2-positive cells in the striatum have increased activity during tadpole begging, suggesting a function for this brain region in tadpole signaling. This is supported by many studies regarding the role of FoxP2 in the striatum of vocalizing birds and mammals. Deficits in songbird vocalizations are observed after FoxP2 knockdown in Area X, a striatal nucleus involved in song learning [24]. At a cellular level, FoxP2 has been implicated in structural plasticity, where FoxP2 modifications influence spiny dynamics of Area X neurons in zebra finches [60] and dendrite lengths of striatal neurons in mice [61]. In this same study, the variant of FoxP2 expressed in these mice also impacted dopamine concentrations in the striatum and nucleus accumbens. Dopamine signaling is critical to tadpole begging behavior [39], and our results here suggest a potential role for FoxP2 in dopamine signaling that should be investigated in the future. This general cellular dysregulation can be seen in mice with FoxP2 mutations, where the striatum is more active and motor-skill learning is disrupted due to abnormal temporal coordination of striatal firing [62]. Together, our work expands the potential role of FoxP2 in the striatum to behavioral signaling in amphibians, suggesting a conserved role for striatal FoxP2 in communication across tetrapod vertebrates.

The cerebellum is a highly conserved vertebrate brain region that coordinates voluntary movements and motor learning [63]. The cerebellum is also implicated in language [64], as there is higher overall cerebellar activity during language tasks in humans [65]. Mice expressing the FoxP2 with the R552H mutation (that leads to speech-language disorders in humans) have impaired ultrasonic vocalizations and poor dendritic development of FoxP2-positive cerebellar Purkinje cells [46]. A reduction of FoxP2 expression specifically in cerebellar Purkinje neurons leads to a reduction of ultrasonic vocalizations in mouse pups [66]. Moreover, expressing the wild type human FOXP2 in the cerebellum partially rescues ultrasonic vocalizations in mice with global expression of FoxP2 with the R552H mutation [67]. To our knowledge, the role of FoxP2 in the cerebellum during vocal learning in songbirds is unknown, but cerebellar lesions impair song learning [68]. Our study, along with studies in neonatal mice, suggest that investigating the role of cerebellar FoxP2 during vocal signaling in songbirds would resolve whether the role of these neurons in coordinating motor signaling are generalizable across taxa.

The nucleus accumbens brain region in the basal ganglia involved in motivation and behavioral reinforcement. In this study, we found increased colocalization of pS6 and FoxP2 in the nucleus accumbens of aggressive tadpoles compared to controls. In mice, increased neural activation in the nucleus accumbens is observed with aggression seeking behavior [69]. Only one study, to our knowledge, has examined the role of FoxP2 specifically in the nucleus accumbens, where deletion in adult mice leads to altered reward and fear learning ([70]; this study did not report effects on aggression). Heterozygous FoxP2^+/-^ mice also show altered aggression in resident-intruder and maternal aggression assays, although the brain regions regulating these altered behaviors was not studied [31]. Our data suggests investigating nucleus accumbens FoxP2 function in the context of aggression would be a fruitful avenue of future research.

### Summary

We present evidence that FoxP2 has conserved brain expression patterns across vertebrates by filling in a critical taxonomic gap from amphibians. We also show that parent-directed signaling by tadpoles is associated with activity of FoxP2-positive cells in the striatum and cerebellum. In contrast, activity of FoxP2-positive cells in the nucleus accumbens was associated with aggression. Overall, this work supports the hypothesis that the FoxP2 transcription factor is part of a molecular toolkit important for social behavior via striatal and cerebellar circuits across many animals.

## Supporting information

analysis code

## Data Accessibility

All data are included in supplementary materials.

## Acknowledgements

We thank Madison Lacey and David Ramirez for their support maintaining our poison frog colony. We especially appreciate comments on early versions of this manuscript from Neil Khosla. We acknowledge that our work takes place on the ancestral and unceded land of the Muwekma Ohlone tribe.

## Author contributions

Conceptualization: SCL and LAO

Methodology: SCL, BCG

Formal Analysis: SCL and LAO

Investigation: SCL, MGR, BCG, AAC, and JEM

Resources: LAO

Data Curation: SCL

Writing – original draft preparation: SCL and LAO

Writing – review and editing: JEM, MGR, and JMB

Visualization: SCL, BCG, AAC

Supervision: LAO, JMB

Project administration: LAO

Funding acquisition: LAO

