## Supplementary Materials for "Activity of FoxP2-positive neurons is associated with tadpole begging behavior"

Sarah C. Ludington, Jordan E. McKinney, Julie M. Butler, Billie C. Goolsby, Ashlyn A. Callan, Maiah Gaines-Richardson, Lauren A. O'Connell

Department of Biology, Stanford University, Stanford, CA 94305, USA

**<u>Tadpole behavior in begging and aggression groups</u>**

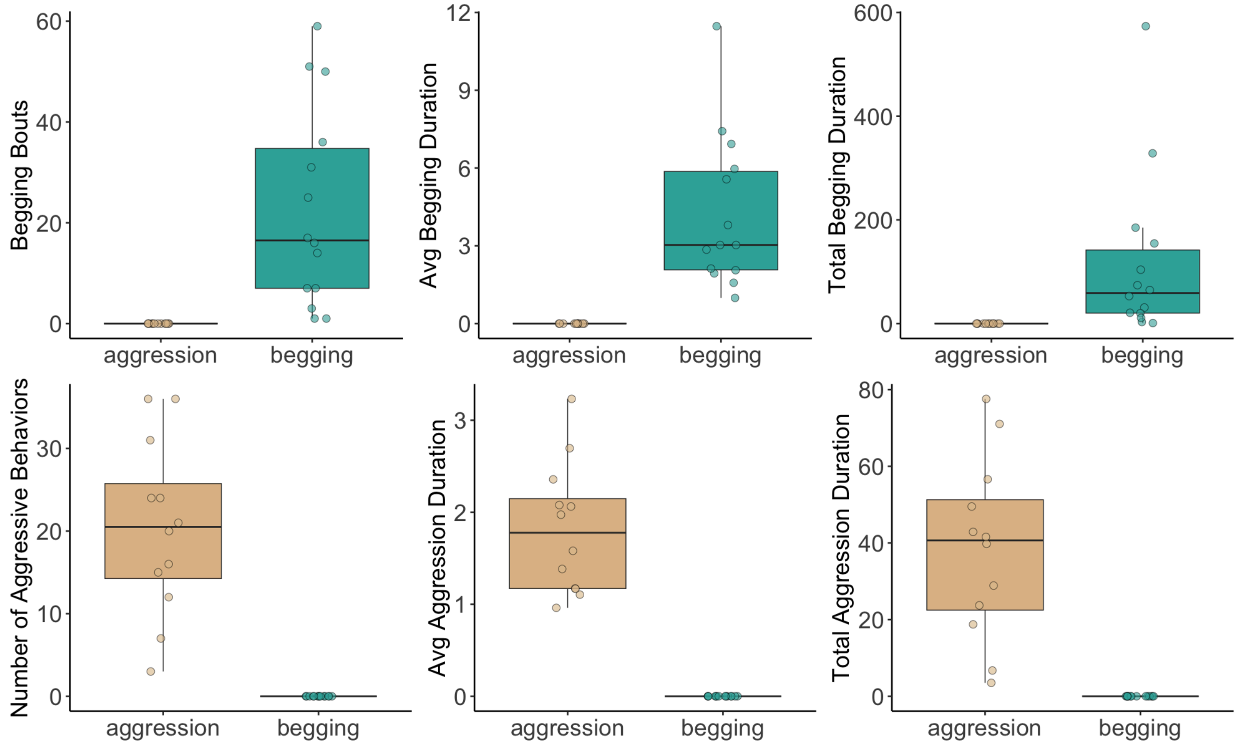


**Figure S1.** Behavior scores are shown for begging (green, top) and aggression (orange, bottom) groups and include the number of behavioral events (left panel), the average duration (middle panel, in seconds), and the total duration (right panel, in seconds).

### Validation of the FoxP2 antibody in the mimic poison frog (*Ranitomeya imitator*)

#### Obtaining FoxP1, FoxP2, and FoxP4 mRNA sequences

*Obtaining cDNA:* We rapidly dissected the brain from a Gosner stage 40 *Ranitomeya imitator* tadpole on ice, which we then lysed in buffer RLT (Qiagen #79216, Germantown, MD, USA) with 1% beta-mercaptoethanol into a Bead Mill 24 homogenizer (ThermoFisher #15-340-163, Waltham, MA, USA). We then transferred homogenate supernatant to a gDNA eliminator spin column and proceeded with RNA isolation using the RNeasy Plus Mini Kit (Qiagen #74134), according to manufacturer instructions. We quantified RNA concentration on a NanoDrop One spectrophotometer (ThermoFisher #ND-ONE-W), after which we diluted to load 1 μg of brain RNA with 4 μL of iSCRIPT reverse transcriptase supermix (Bio-Rad #1708840, Hercules, CA, USA) for a final volume of 20 μL. We produced cDNA using the following thermal cycler parameters (25°C,5:00; 46°C, 30:00; 95°C, 1:00; C1000 Thermal Cycler, Bio-Rad #1851148). We quantified and recorded cDNA quantity on the nanodrop.

*PCR validation of* foxp2*:* We retrieved the *foxp1*, *foxp2*, and *foxp4* sequences (GenBank accession numbers: PP952377-9) from our in-house transcriptome to be published at a later date. We next sought to validate the *foxp2* sequence in order to design probes for RNAscope detection and visualization of RNA in tissue sections. We designed the custom primers (Integrated DNA Technologies, San Diego, CA, USA) to amplify the *R. imitator foxp2* sequence with a T3 polymerase recognition site added to the reverse sequence (Forward: 5'CAAAGCATCACCACCAATAA; Reverse: 5'aattaaccctcactaaagggGTAAGCAAACGTCCGTGTA). Using the cDNA as a template, we ran a PCR with 12.5 μL of platinum PCR hot start master mix, 0.5 μL 10 μM forward primer, 0.5 μL 10 μM reverse primer, 10.5 μL nuclease free water, and 1μL cDNA for a total volume of 25 μL which we incubated with the following thermal cycler parameters (94°C,2:00; 94°C, 0:30; 55°C, 0:30; 72 1:00; repeat 45 cycles). We evaluated PCR products on a 1% agarose gel in 1x TAE buffer with 0.01% GelGreen nucleic acid gel stain (Biotum #41005, San Francisco, CA, USA). We loaded 3 μL of GeneRuler 1kb Plus DNA ladder (ThermoFisher #SM1331) and 1 μL of 6X gel loading dye (ThermoFisher #R1161 + 2 μL of PCR in the experimental well. We ran the gel at 135 V for 20 minutes. After confirming presence of PCR product at the correct size (~700 bases), we purified the product using manufacturer instructions for the MinElute PCR Kit (Qiagen #28004) and used the forward primer for Sanger Sequencing using GeneWiz (San Francisco, CA, USA). Our Sanger sequence results confirmed that our purified product was that of *foxp2*. Following sequence confirmation, we provided the sequence to Advanced Cell Diagnostics (Newark, CA, USA) which designed a new 20ZZ probe (#NPR-0052782) targeting the coding sequence in length positions 731-1893, which also aligns to the FOXP2 sequence of the Sardinian Tree Frog (*Hyla sarda,* NCBI Reference Sequence: XM_056573878.1).

#### Comparison of immunogen sequence of FOXP2 antibody to other species

The immunogen used to make the FoxP2 antibody used in this study was a synthetic peptide corresponding to the Human FOXP2 amino acids 703-715 (C-terminal). The human-based sequence of the synthetic peptide is C-REIEEEPLSEDLE. We compared the sequence of the synthetic peptide with several species, including *R. imitator*, to evaluate whether the antibody would bind frog FoxP2. We found this segment of the *R. imitator* FoxP2 peptide sequence to be the same as *Xenopus laevis* and one amino acid different from mammals and birds, with an overall similarity of roughly 92% to human FOXP2. In contrast the epitope is only 54% and 31% similar to *R. imitator* FoxP1 and FoxP4, respectively (Figure S2).

| | | | | | | | | | | | | | |
|---|---|---|---|---|---|---|---|---|---|---|---|---|---|
| Antibody Epitope | R | E | I | E | E | E | P | L | S | E | D | L | E |
| Human FOXP2 | R | E | I | E | E | E | P | L | S | E | D | L | E |
| Mouse FoxP2 | R | E | I | E | E | E | P | L | S | E | D | L | E |
| Zebra finch FoxP2 | R | E | I | E | E | E | P | L | S | E | D | L | E |
| *Xenopus* FoxP2 | R | E | L | E | E | E | P | L | S | E | D | L | E |
| *R. imitator* FoxP2 | R | E | L | E | E | E | P | L | S | E | D | L | E |
| *R. imitator* FoxP1 | R | D | Y | E | D | E | P | V | N | E | D | M | E |
| *R. imitator* FoxP4 | H | D | L | D | P | E | T | A | M | E | D | L | S |

**Figure S2. Sequence comparison of the FoxP2 antibody immunogen.** Sequences from mouse (AAH58960.1), zebra finch (NP_001041728.1), and *Xenopus* (NP_001089138.1) were obtained from Genbank. The *R. imitator* sequence FoxP2 was obtained from the genome (Genbank accession: GCA_032444005.1). The *R. imitator* FoxP1 and FoxP2 sequences were obtained from an in-house transcriptome. All three *R. imitator* FoxP sequences are available on GenBank (Accession PP952377-9).

**No primary antibody control**

Our immunohistochemistry experiments included no primary control, which blocked all signal (Figure S3).

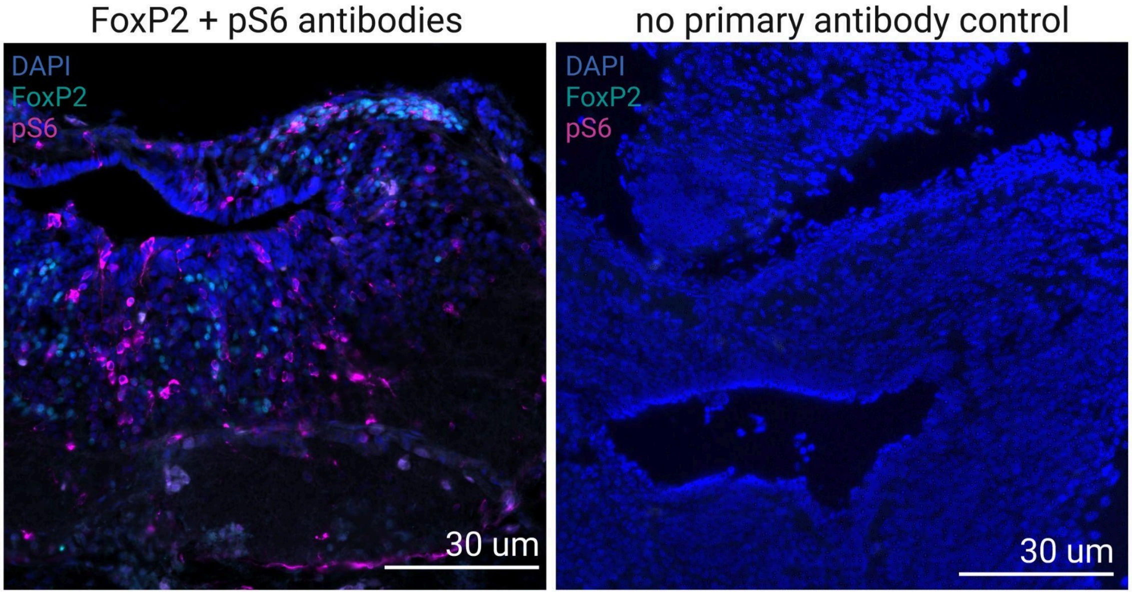


**Figure S3. Negative control slides testing FoxP2 antibody specificity.** Representative micrographs of hindbrain brain structures stained with either FoxP2 and pS6 primary antibodies (left) or a no primary antibody control (right). Control slides were incubated with blocking solution only. Composite images show DAPI (blue), FoxP2 (cyan), and pS6 (pink). Scale bars at 30 μm.

**Western Blot**

*Western methods*: To further test antibody specificity, we performed a western blot for FoxP2 using the same primary antibody used in immunohistochemistry (Figure S3), but a different anti-goat secondary antibody suitable for western blot (i.e. with a peroxidase tag). Protein was extracted from three brains by homogenizing the pooled brains in 300 µl lysis buffer (2% SDS + 1x protease inhibitor) and centrifuging for 90 min at 4C. We loaded 40 µg of protein, Dual Color Precision Plus Protein Standards (BioRad 1610374), and a no-protein control (protein ladder, and control in duplicate) into a 12% Mini-PROTEAN TGX Precast gel (BioRad 4561045), and ran the gel for 40 min at 200V. Protein was transferred to a 0.2 µm nitrocellulose membrane using the Trans-Blot Turbo Transfer system (transfer packs: BioRad 1704158) with the Mini-TGX protocol (2.5A constant, up to 25V) for 4 minutes. The membrane was blocked in 1X tris buffered saline with 0.1% tween-20 (TBST) and 5% normal donkey serum on a nutator overnight at 4C. All remaining steps were done at room temperature on a nutator. The membrane was cut in half, with one half left in blocking solution and the other placed in primary antibody solution (anti-FOXP2 at 1:500 in blocking solution) for 1.5 hours. After washing with 1x TBST, the membranes were incubated in bovine anti-goat IgG HRP (Jackson labs #805-035-180) prepared in a blocking solution for 2 hours. After washing, the membranes were reacted with Opti-4CN (BioRad 1708235) for 30 minutes to visualize protein staining. The membranes were then rinsed with deionized water and placed on an LED lightpad for visualization and imaging.

*Western results:* We tried multiple different anti-goat secondaries suitable for western blotting, but all resulted in some non-specific binding despite adjusting blocking and incubation conditions. This non-specific binding is not present with the anti-goat secondary used in immunohistochemistry, as evidenced by the lack of non-specific staining when the primary antibody was omitted from the staining (See Figure S3). The nonspecific binding in the western produced a series of bands between 40-60 kD in both the no-primary and anti-FOXP2 lanes, suggesting this background binding is due to non-specific binding of the secondary antibody (Figure S4). Only the membrane incubated in the FOXP2 primary antibody had bands present between 75-100 kD, consistent with the predicted weight of FoxP2 at ~80 kD, which were not present in the no-primary control. Two faint bands are present in this size range. This could be the FOXP2 antibody binding multiple FoxP2 isoforms with similar molecular weights, as seen in other animals [1–3]. Alternatively, the FoxP2 antibody could be binding to other FoxP members, such as FoxP1 or FoxP4. We consider this unlikely, as the epitope is 92% similar to *R. imitator* FoxP2 and only 54% and 31% similar to *R. imitator* FoxP1 and FoxP4, respectively, although expression of the proteins in cell culture would be required to validate this assumption.

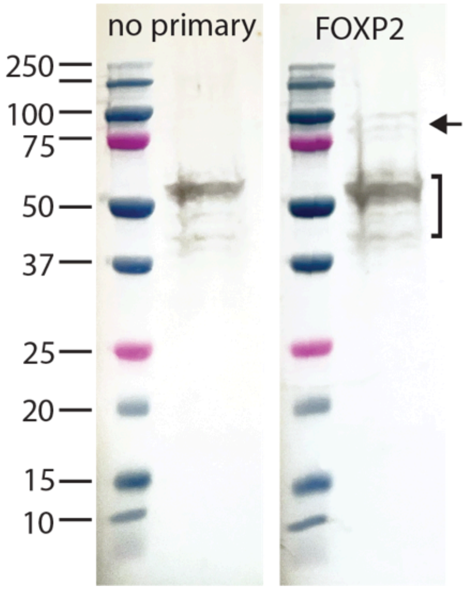


**Figure S4. Western blot using a FoxP2 antibody and protein from *Ranitomeya imitator* tadpole brains.** Omitting the primary FoxP2 antibody resulted in non-specific binding (40-60 kD shown with brackets), as seen on the left blot, which indicates background from the secondary antibody. Incubation with FOXP2 primary antibody at 1:500 (blot on the right) produced two faint bands between 75-100 kD (arrow), consistent with the predicted weight of ~80 kD FoxP2.

**Double labeling of FOXP2 (antibody) and *foxp2* (mRNA)**

*Double labeling methods*: To validate whether the FOXP2 antibody labeled FoxP2-positive cells in *R. imitator*, we sacrificed a stage 30 *R. imitator* tadpole in RNAse free 1X Phosphate Buffered Saline (PBS) with 20% benzocaine, euthanized by rapid cervical dislocation, and then embedded the head in Tissue-Tek OCT embedding medium (Sakura #4583, Torrance, CA, USA), which we then froze on dry ice. We sectioned the brain at -18°C at 15 µm into four series on Superfrost Plus slides (VWR #48311-703, Visalia, CA, USA). After sectioning was completed, slides were stored at -80C until processing. One series of sections was used for RNAscope *foxp2* mRNA transcript labeling. We used the RNAscope multiplex fluorescent V2 assay (ACD #323270), using manufacturer instructions for fresh frozen tissues (ACD Document #UM323100). We used the fluorescent dye Opal 570 (dilution 1:1500 in tyramide signal amplification buffer, Akoya Biosciences SKU#FP1488001KT, Menlo Park, CA, USA). A second series of sections was used for RNAscope + immunohistochemistry, which was performed using manufacturer instructions for in situ hybridization (ISH)/immunohistochemistry (IHC) (ACD Technical Note #323100-TN). Briefly, after completing the HRP blocking step in RNAscope, we washed slides in TBST (0.05% of 1XTBS with 10% Tween 20) twice for 2 minutes each, performed blocking with 10% normal goat serum in TBS-1% BSA for two hours at room temperature, then performed overnight primary blocking of 1:500 primary antibody in TBS-1% BSA at room temperature. The next day, we washed the slides in TBST twice for 5 minutes each. We then incubated the slides with HRP-conjugated secondary antibody (1:1000, Peroxidase AffiniPure Donkey Anti-Goat, Jackson Immunoresearch #705035003, West Grove, PA, USA) in TBS-1% BSA for 30 minutes at RT, washed with TBST for 5 minutes at room temperature twice. Then, slides were treated with Opal 570 (1:750) for 10 minutes at RT. Slides were washed twice with TBST for 2 minutes at room temperature twice, then stained with DAPI and mounted. An additional series of sections was used for immunohistochemistry following the established protocol reported in the main manuscript. By comparing the A, B, and C series we can compare staining quantities across relatively identical sections (15 µm-30 µm distance).

*Double labeling results:* We compared where staining occurred across a tadpole brain. We first costained one series together (Figure S5A), which caused weak antibody labeling, which we theorize is due to the protease component involved in RNAscope labeling. Nonetheless, we identified costained FoxP2-positive cells using both RNAscope and immunohistochemistry in the same tissue. We then used sections representative of the same brain region in the same animal in alternate series to separate immunohistochemistry and RNAscope to overcome any protocol incompatibilities. We found nuclear staining of FOXP2-positive cells using immunohistochemistry and puncta staining of *foxp2* transcripts using RNAscope (Figure S5B-E).

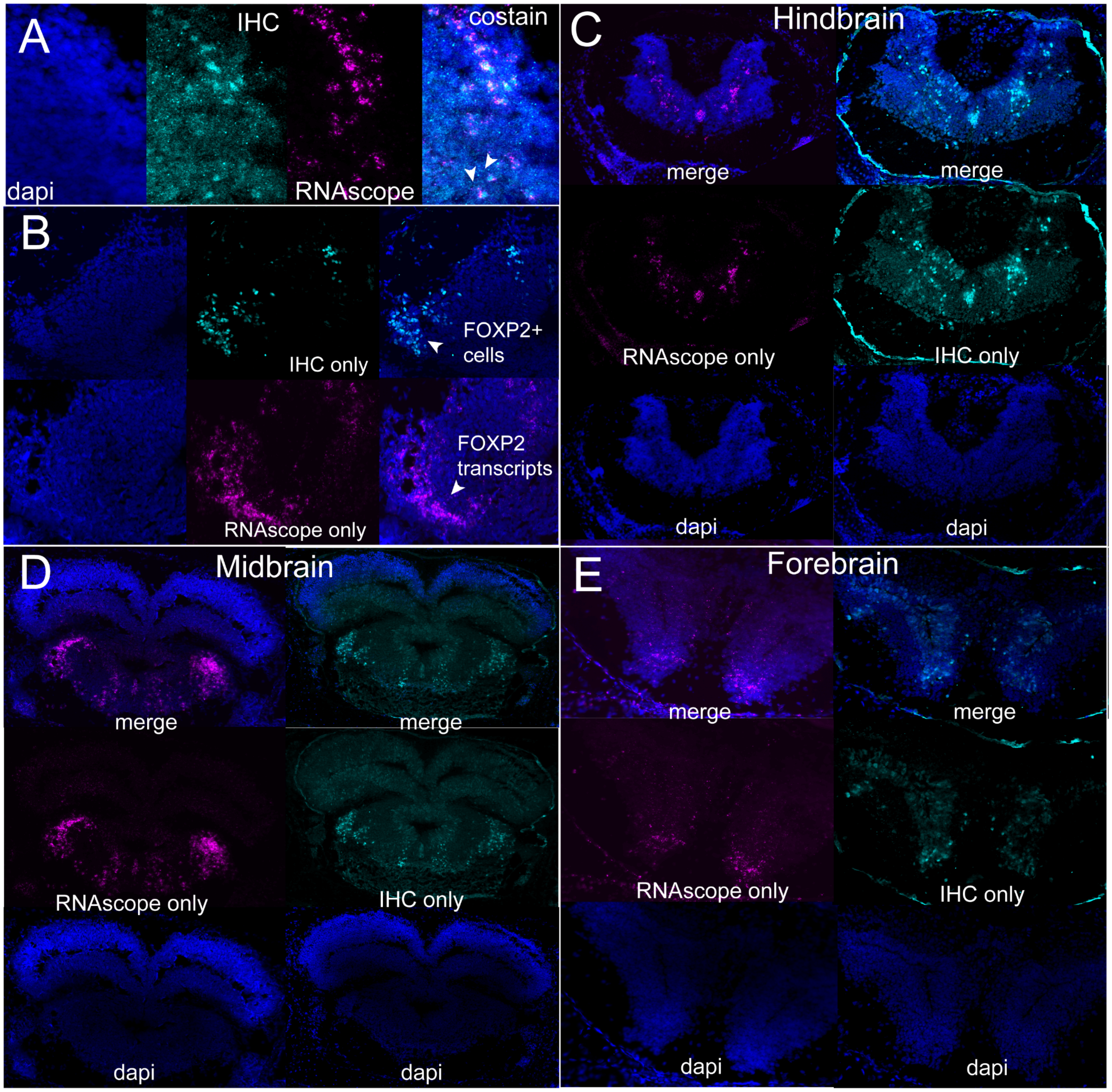


**Figure S5. Comparison of RNAscope labeling of *foxp2* transcripts and immunohistochemical labeling of FoxP2-positive neurons.** We compared where staining occurred across a tadpole brain. We first **(A)** costained one series together, which caused a high background in antibody labeling, which we theorize is due to the protease component involved in RNAscope labeling. Nonetheless, we identified costained FoxP2-positive cells using both RNAscope and immunohistochemistry in the same tissue. We then (**B)** used sections representative of the same brain region in the same animal in alternate series to separate immunohistochemistry and RNAscope to overcome any protocol incompatibilities. We found nuclear staining of FoxP2-positive cells using immunohistochemistry and puncta staining of *foxp2* transcripts using RNAscope in the same brain region. We highlight similar staining patterns in the **(C)** hindbrain around the Gc and Cb, the **(D)** midbrain around the Av, and **(E)** the forebrain in the striatum.

### Brain distribution of FoxP2-positive cells

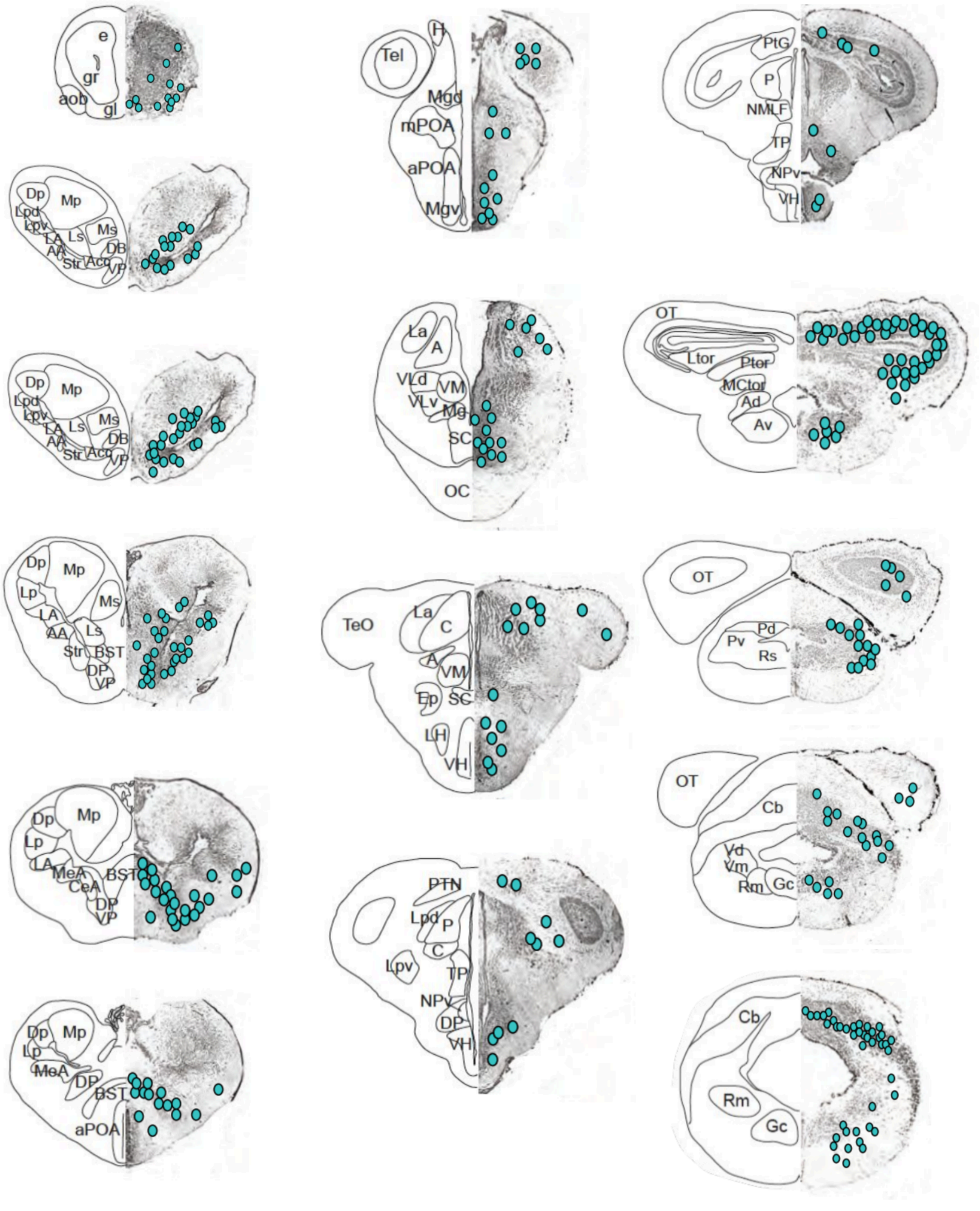


**Figure S6. Neural distribution of FoxP2 in the tadpole brain.** FOXP2 is widely distributed throughout the *R. imitator* tadpoles brain. Abbreviations: A, anterior thalamic nucleus; AA, anterior amygdaloid area, NAcc, nucleus accumbens; Ad, anterodorsal tegmental nucleus; AH, anterior hypothalamus; aob, accessory olfactory bulb; Av, anteroventral tegmental nucleus; BST, bed nucleus of the stria terminalis, C, central thalamic nucleus, Cb, cerebellum; CeA, central amygdala; DB, diagonal band of Broca; DH, dorsal hypothalamic nucleus; Dp, dorsal pallium;

DP, dorsal pallidum; e, postolfactory eminence; Ep, posterior entopenducular nucleus; Gc, griseum central rhombencephali; gl, glomerular layer of the olfactory bulb; gr, granule cell layer of the olfactory bulb; Hv, ventral benula; La, lateral thalamic nucleus, anterior division; LA, lateral amygdale; LH, lateral hypothalamic nucleus; Lp, lateral pallium; Lpd, lateral thalamic nucleus, posterodorsale; Lpv, lateral thalamic nucleus, posteroventrale; Ls, lateral septum; M, dorsal midline; MeA, medial amygdale; Mgd, dorsal magnocellular nucleus, dorsal part; Mgv, ventral magnocellular nucleus, ventral part; ml, mitral cell layer of the olfactory bulb; Mp, medial pallium, Ms, medial septum; ON, optical nerve; Npv, nucleus of the periventricular organ; P, posterior thalamic nucleus; Pd, nucleus posterodorsalis tegmenti; aPOA, anterior preoptic area; mPOA, medial preoptic area; Pv, nucleus posteroventralis tegmenti; Rm, nucleus reticulris medius; Rs, nucleus reticularis superior; SC, suprachiasmatic nucleus; Str, striatum; Tect, optical tectum; Tel, telencephalon; Tor-L, torus semicirularis, laminar nucleus; Tor-P, torus semicircularis, principal nucleus; Tor-V, torus semicuricularis, ventral area; TP, posterior tuberculum; Vd, descending trigeminal tract; VH, ventral hypothalamic nucleus; VLd, ventrolateral thalamic nucleus, dorsal part; VLv, ventrolateral thalamic nucleus, ventral part; Vm, nucleus motorius nervi trigemini; VM, ventromedial thalamic nucleus; VP, ventral pallidum.

**Number of FoxP2-positive cells across brain regions**

The number of FoxP2-positive cells varied between behavioral group and brain region (group*region: $F_4 = 28.644$, $p < 0.001$) (Figure S7). Aggressive tadpoles had less FoxP2-positive cells than begging and control tadpoles in the striatum (Str, aggression vs begging: $z = -2.667$, $p = 0.012$; aggression vs control: $z = -3.762$, $p < 0.001$). Groups did not differ in the number of FoxP2-positive cells in the nucleus accumbens (NAcc) or cerebellum (Cb).

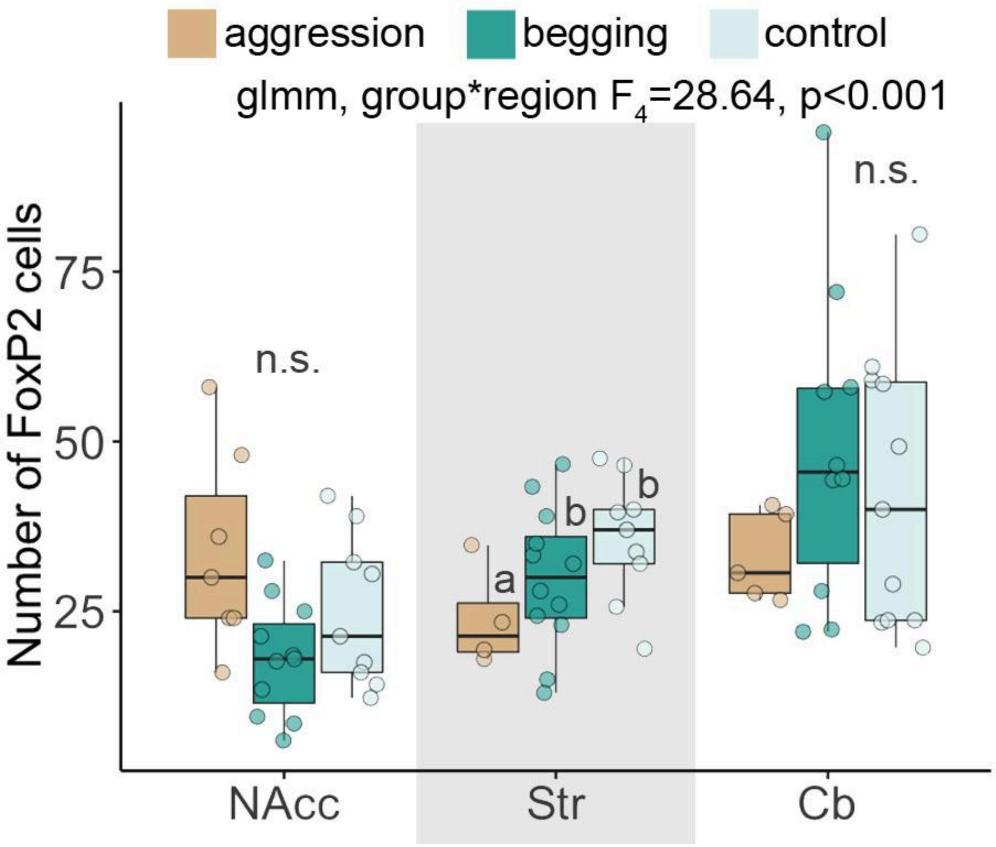

**Figure S7. Number of FoxP2 cells across brain regions and behavioral groups.** The number of FoxP2-positive cells in aggressive (orange), begging (dark green) and control (light green) tadpoles are shown in boxplots with individual tadpoles displayed in dots. Groups not connected by the same letter are significantly different. Abbreviations: NAcc, nucleus accumbens; Str, striatum; Cb, cerebellum.

### Correlations of neural activity with behavior

As the proportion of FoxP2-positive cells that are also pS6-positive was higher in the striatum and cerebellum of begging tadpoles, we asked whether FoxP2-positive cell activity was correlated with begging behavior (Figure S8). We examined the correlation between the number of active FoxP2-positive cells in the striatum and cerebellum with the number of begging events, the average begging duration, and the total begging duration. No significant correlation was found in any comparison.

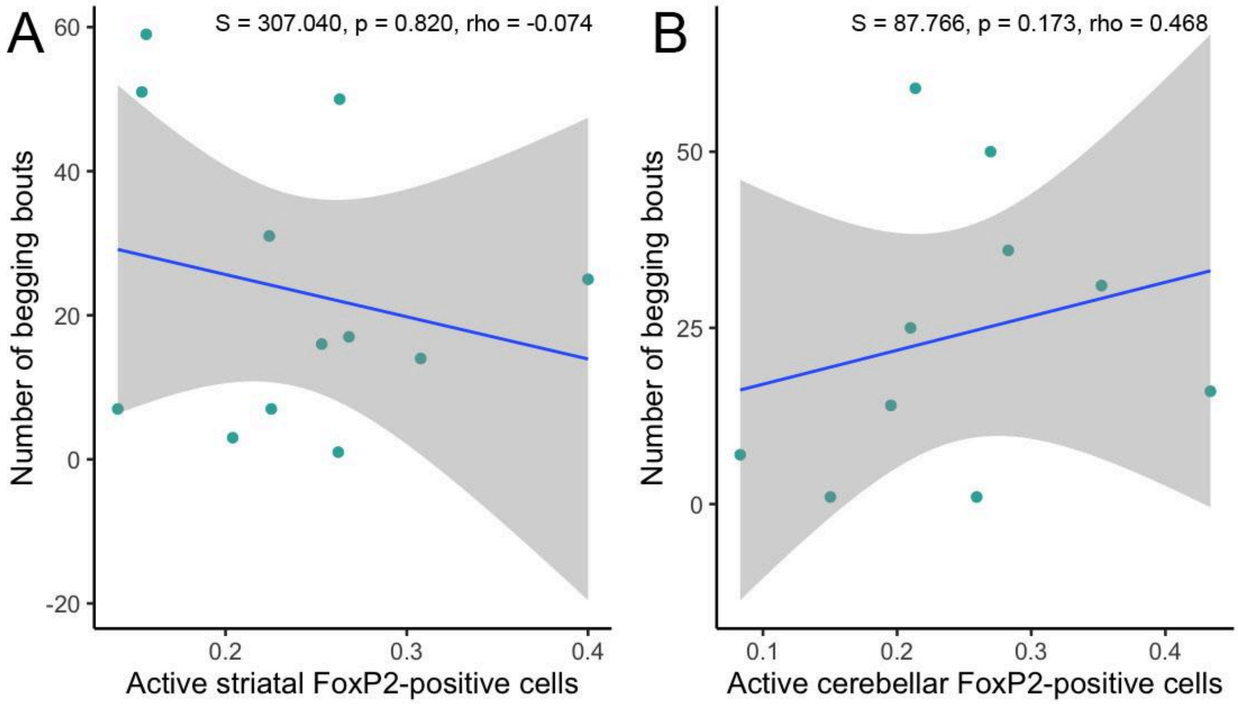


**Figure S8. Begging behavior does not correlate with activity of FoxP2-positive cells.** The number of tadpole begging bouts does not correlate with the proportion of FoxP2-positive cells that are pS6-positive in the **(A)** striatum or **(B)** cerebellum.
